# Batch experiments demonstrating a two-stage bacterial process coupling methanotrophic and heterotrophic bacteria for 1-alkene production from methane

**DOI:** 10.1101/2021.08.17.456502

**Authors:** Ramita Khanongnuch, Rahul Mangayil, Ville Santala, Anne Grethe Hestnes, Mette Marianne Svenning, Antti J Rissanen

## Abstract

Methane (CH_4_) is a sustainable carbon feedstock source for aerobic CH_4_-oxidizing bacteria (methanotrophs) to produce value-added chemicals. Under substrate-limited (e.g., CH_4_, oxygen and nitrogen) conditions, CH_4_ oxidation results in the production of various short-chain organic acids and platform chemicals. These CH_4_-derived products could be broadened by utilizing them as a feedstock for heterotrophic bacteria. As a proof of concept, a two-stage system for CH_4_ abatement and 1-alkene production was developed in this study. Types I and II methanotrophs, i.e., *Methylobacter tundripaludum* SV96 and *Methylocystis rosea* SV97, respectively, were investigated in batch tests under different CH_4_ and air supplementation schemes. CH_4_ oxidation under either microaerobic or aerobic conditions induced the production of formate, acetate, succinate, and malate in *M. tundripaludum* SV96, accounting for 4.8-7.0% of consumed CH_4_-carbon while *M. rosea* SV97 produced the same compounds except for malate, and with lower efficiency than *M. tundripaludum* SV96, accounting for 0.7-1.8% of consumed CH_4_-carbon For the first time, the organic acids-rich spent media of methanotrophs were successfully used for 1-alkene production using engineered *Acinetobacter baylyi* ADP1 ‘*tesA-undA* cells. The highest yield of 1-undecene was obtained from spent medium of *M. tundripaludum* SV96 at 68.9 ± 11.6 μmol mol C_substrate_^−1^.

## 1. Introduction

Methane (CH_4_) is the second most important greenhouse gas (GHG) after CO_2_, having a global warming potential (GWP) of approximately 30 times higher than CO_2_ on a 100-year time horizon (IPCC, 2021). Its emission from anthropogenic activities has been continuously increasing, being approximately as high as 60% of total CH_4_ emission to the atmosphere (based on top-down estimates of Saunois et al., 2020). Hence, keeping on a stringent climate policy for the next several decades, particularly the use of CH_4_ as an energy source, would significantly reduce CH_4_ emissions (Harmsen et al., 2020). In environmental carbon flux, CH_4_ oxidizing bacteria are the key regulators for CH_4_ abatement (Hanson and Hanson, 1996; Saarela et al., 2020). Due to its abundance and potentiality as a sustainable carbon feedstock, the development of the biological processes for CH_4_ conversion to bio-based chemicals/liquid fuel is promising and attractive research area (Liu et al., 2020; López et al., 2013; Sun et al., 2018).

Aerobic methane oxidizing bacteria (methanotrophs) containing methane monooxygenases (MMO) use CH_4_ as the sole carbon and energy source and oxygen as an electron acceptor. Methanotrophs are classified into gammaproteobacterial (Types I and X) and alphaproteobacterial (Type II) methanotrophs based on their different pathways for formaldehyde assimilation into biomass, i.e., the ribulose monophosphate (RuMP) and the serine cycles, respectively (Kalyuzhnaya et al., 2015). Along with these pathways, CH_4_ is potentially converted into various value-added products including methanol, single cell protein, ectoine and soluble metabolites (Cantera et al., 2018; Ge et al., 2014; Sheets et al., 2016; Strong et al., 2016). In various environmental processes and biological systems, methanotrophs have been reported to support other bacteria by producing organic carbon sources from CH_4_, such as in wastewater treatment (Cao et al., 2019; Costa et al., 2000) and heavy metal bioremediation (Lai et al., 2016). The pathways for the excretion of organic acids (e.g., formate, acetate, lactate and succinate) have been observed in the experiments and deciphered from genomes of Type I methanotrophs during O_2_ limiting conditions including the Embden-Meyerhof-Parnas (EMP) and tricarboxylic acid (TCA) cycle (Fig. 1) (Gilman et al., 2017; Kalyuzhnaya et al., 2013). Besides O_2_ limiting conditions, acetate production was also very recently detected in liquid culture of methanotrophs incubated under aerobic conditions (Lee et al., 2021; Takeuchi and Yoshioka, 2021). These studies also showed the potential of the methane-derived organic acids in biotechnological applications by cultivating Type I methanotrophs and heterotrophs in co-culture systems, where the heterotrophs utilized the organic acids produced by the methanotrophs (Lee et al., 2021; Takeuchi and Yoshioka, 2021). For example, acetate produced by *Methylocaldum marinum* S8 was successful used as a growth medium for *Cupriavidus necator* cultivation (Takeuchi and Yoshioka, 2021) and co-culture of *Methylococcus capsulatus* Bath and the engineered *Escherichia coli* SBA01 was carried out for mevalonate production (Lee et al., 2021).

**Fig. 1.**
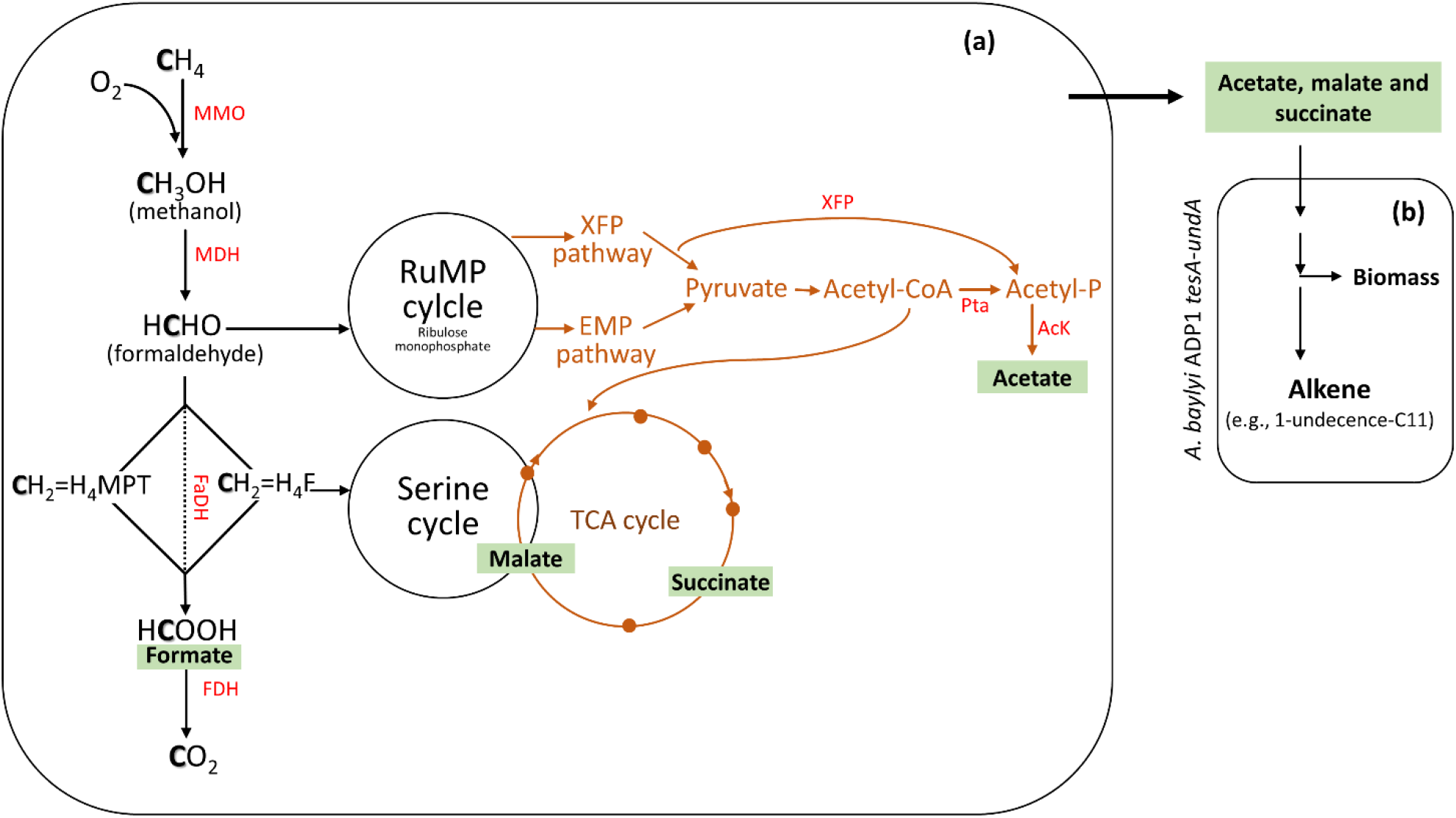
The principle of a two-stage bacterial process. The estimated pathway of the production of various organic acids could occur in methanotrophs (a). Organic acid rich spent medium was fed to the engineered *Acinetobacter baylyi* ADP1 for alkene production (b). Aerobic CH_4_ oxidation pathway in Type I (RuMP cycle) and Type II (Serine cycle) methanotrophs (black arrows) and the possible pathway of CH_4_ oxidation under O_2_-limiting conditions via glycolysis-based CH_4_ fermentation mode (brown line). Abbreviations: **RuMP**, ribulose monophosphate; **EMP**, Embden–Meyerhof–Parnas; **TCA**, Tricarboxylic acid; **MMO**, methane monooxygenase; **MDH**, methanol dehydrogenase; **FaDH**, formaldehyde dehydrogenase; **H4F**, tetrahydrofolate pathway; **H4MPT**, tetrahydromethanopterin pathway; **FDH**, formate dehydrogenase; **PTa**, phosphate acetyl-transferase; **AcK**, acetate kinase; **XFP**, phophoketolase xylulose 5-phosphate/fructose 6-phosphate phosphoketolase.

*Acinetobacter baylyi* ADP1, naturally competent gram-negative gammaproteobacteria, have been recently gained interest in bioengineering. The bacterium’s capacity to metabolize a wide range of carbon sources and genetic tractability has led to its application in long-chain alkyl ester and 1-alkene compounds (e.g. 1-undecene) biosynthesis from aromatic lignin-derived monomers and lignocellulose (Luo et al., 2019; Salmela et al., 2020). As a platform chemical, 1-undecene can be used in the chemical synthesis of medium-chain poly-α-olefins (C33) which are commonly used as lubricants (Salmela et al., 2020). Recently, the ability of *A. baylyi* ADP1 to metabolize fermentation by-products has been exploited to broaden the metabolic landscape in biological production processes (Mangayil et al., 2019; Salmela et al., 2018). Hence, *A. baylyi* ADP1 would provide a promising, testable candidate for the bioconversion of CH_4_–derived organic acids. However, it is critical to find a suitable methanotroph partner for such application. For instance, whether Type I or II methanotroph would make a good candidate for converting CH_4_ to organic acids for *A. baylyi* ADP1 is not known, as to our knowledge, the comparison of organic acid production yields between Types I and II methanotrophs has not been previously reported. Nevertheless, the CH_4_-derived compounds are limited to short carbon chain carbon compounds (C2-C6), which could be further used as a feedstock for various heterotrophic bacteria, that can be easily engineered in order to extend the range of CH_4_-derived value-added (platform) chemicals.

With the aim to redirect the ‘overflow carbon’ from microaerobic methanotrophic fermentation, organic acids obtained from CH_4_ oxidation process could potentially provide as important feedstocks towards high-value product formation. Thus, a two-stage bioprocess setup was designed for successful integration of production of CH_4_-derived organic acids with aerobic synthesis. The growth and metabolite production profiles of Types I and II methanotrophs, i.e., *Methylobacter tundripaludum* SV96 and *Methylocystis rosea* SV97, respectively, were investigated under different gas supplementation schemes. Next, the possibility of using the spent media of the methanotrophs as a growth substrate for the cultivation of wild type *A. baylyi* ADP1 was tested. Finally, the synthesis of 1-alkenes from CH_4_ was demonstrated by a two-stage process with a methanotroph, i.e., *M. tundripaludum* SV96 and *M. rosea* SV97, and an engineered *A. baylyi* ADP1 (‘*tesA*-*undA*) strain (Luo et al., 2019).

## 2. Materials and Methods

### 2.1. Strains and cultivation conditions

*M. tundripaludum* SV96 and *M. rosea* SV97, isolated from Arctic wetland soil, Norway (Wartiainen et al., 2006a, 2006b), were used in this study. Nitrate mineral salts (NMS) medium used for cultivating the methanotrophs was DSMZ 921 medium (German Collection of Microorganisms and Cell Cultures, 2007) with initial pH ~6.80 and modification of adding with 1 μM lanthanum chloride (LaCl_3_). The pre-inoculum was grown in 120-mL serum bottles containing 10 mL NMS medium and 20% CH_4_ and 80% air in headspace and incubated statically at 20 °C. All experiments were conducted under sterile conditions and the serum bottles used for methanotrophs cultivation were sealed with butyl rubber stoppers and capped with aluminum crimps.

For cocultivation studies, wild type and the engineered *A. baylyi* ADP1 strain (Luo et al., 2019) carrying the plasmid pBAV1C-*‘tesA*-*undA* (*A. baylyi* ADP1 ‘*tesA-undA*) were used. For pre-inoculum, *A. baylyi* ADP1 cells were inoculated in 10 mL culture tubes containing LB medium (5 g L^−1^ yeast extract, 10 g L^−1^ tryptone and 5 g L^−1^ NaCl) supplemented with 0.5% glucose and 25 μg mL^−1^ chloramphenicol (for *A. baylyi* ADP1 ‘*tesA-undA*). The inoculated tubes were aerobically grown overnight at 30 °C and 300 rpm. For 1-undecene synthesis, *A. baylyi* ADP1 ‘*tesA-undA* was induced with 0.5 mM cyclohexanone (Luo et al., 2019).

### 2.2. Evaluation of organic acid production by *M. tundripaludum* SV96 and *M. rosea* SV97

Batch tests were performed in 120-mL airtight serum bottles with a working volume of 15 mL NMS medium. The precultures of *M. tundripaludum* SV96 and *M. rosea* SV97 were inoculated with an initial optical density at 600 nm (OD_600nm_) of 0.02 and the growth, CH_4_ utilization and organic acid profiles were monitored every two days for 14 days at 20°C and static conditions. Both methanotrophs were tested under three different gas supplementation schemes (Fig. 2). On day 0, the bottles were filled with 20% CH_4_ and 80% air into headspace accounting for the initial O_2_/CH_4_ molar ratio of ~1.2 and incubated for 7 days. After CH_4_ and O_2_ concentrations were depleted on day 7, the batch bottles were supplemented with three different gas compositions into headspace, i.e., test I: CH_4_ + air (20% CH_4_ and 80% air), test II: only CH_4_ (20% CH_4_ and 80% N_2_), and test III: only air (20% air and 80% N_2_). The bottles containing only NMS without bacterial cells were used as the control. The bottles were incubated further for 7 days (days 8-14). The biomass growth (OD_600_), gas compositions in headspace and organic acid accumulation in liquid medium were monitored every two days.

**Fig. 2.**
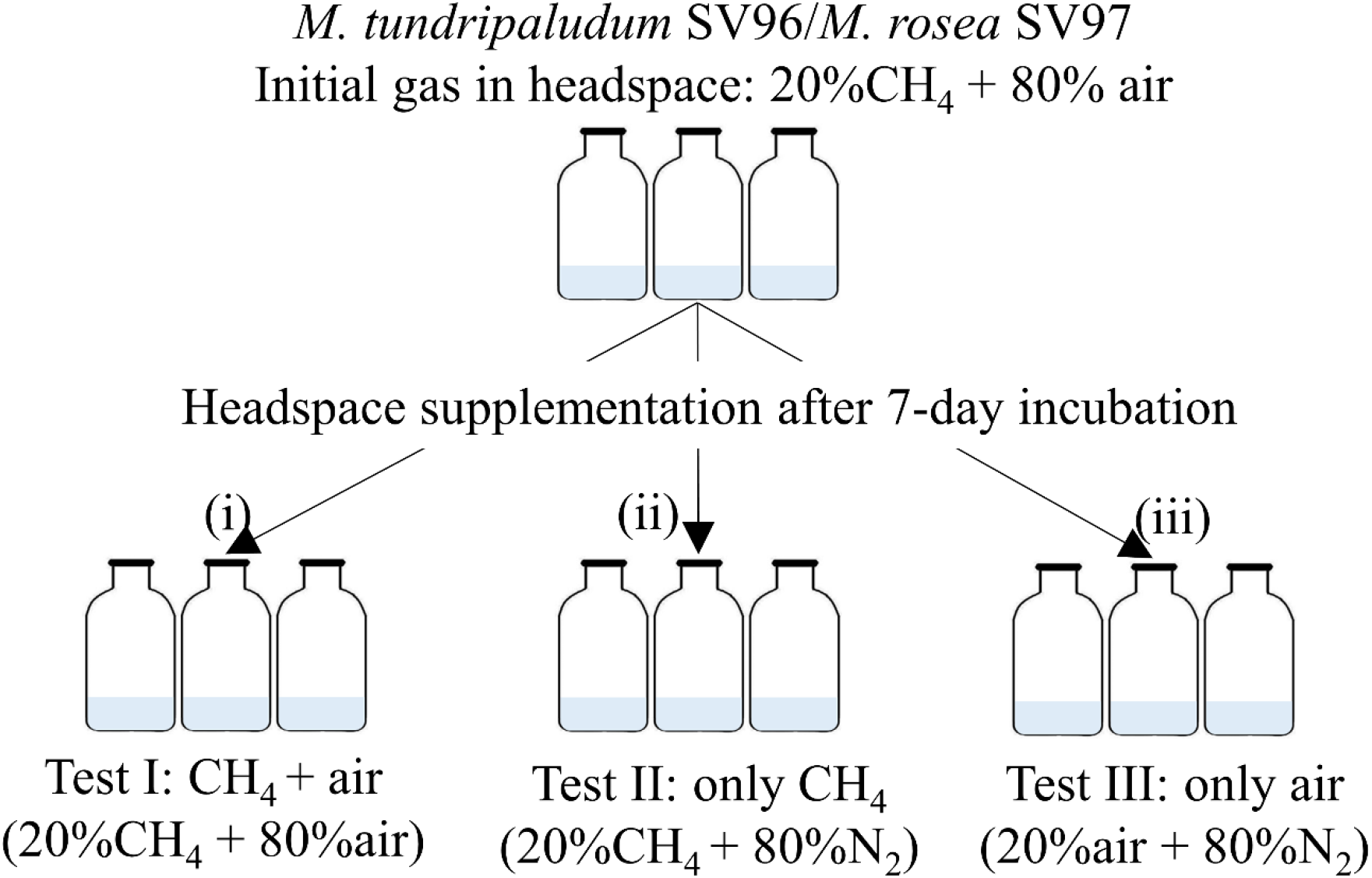
Experimental set up for methanotrophs under three different gas supplementation schemes applied on day 7 including test I: both CH_4_ and air added, test II only CH_4_ added and test III only air added.

### 2.3. Cultivation of wild type *A. baylyi* ADP1 in spent media of different methanotrophs

Prior to the study of 1-alkene production, the capacity of *A. baylyi* ADP1 cells to utilize the spent media of methanotrophs were tested. In this test, *M. tundripaludum* SV96 and *M. rosea* SV97 were cultivated in 500-mL airtight bottles with 60 mL NMS medium in conditions as described previously (Section 2.2). To maximize the organic acid titer, the bottle headspace was initially filled with 20% CH_4_ and 80% air and cultivated for 28 days at optimal conditions for *M. tundripaludum* SV96 and *M. rosea* SV97, individually. Gaseous composition in headspace and organic acid concentrations in liquid medium were monitored twice a week.

After 28 days, the cultivation of methanotrophs was stopped and their spent media were used for *A. baylyi* ADP1 cultivation. The spent media were collected by centrifugation at 12000 rpm for 10 min. The 50-mL supernatant of both methanotrophs were transferred to 250 mL sterile flasks and used for *A. baylyi* ADP1 cultivation. The tests were conducted in duplicate. After 4 h cultivation, the liquid culture was collected to monitor cell growth, organic acid concentration and wax ester production. The spent media of methanotrophs without *A. baylyi* ADP1 were used as contamination control (control 1) and *A. baylyi* ADP1 cells in NMS (fresh medium) were used to determine their background growth (control 2).

### 2.4. Cultivation of *A. baylyi* ADP1 *‘tesA*-*undA* strain and 1-alkene synthesis in spent media of different methanotrophs

In this test, the liquid culture samples from *M. tundripaludum* SV96 and *M. rosea* SV97 microaerobic fermentation were collected and directly used to cultivate *A. baylyi* ADP1 *‘tesA*-*undA* strain. The tests were conducted in quadruplicate in a sealed 20 mL glass tube containing 5 mL of the spent medium. Subsequently, *A. baylyi* ADP1 ‘*tesA*-*undA* cells (initial OD_600nm_, 0.02) were added into the vials. After 1 h of incubation at 30 °C and 300 rpm, 0.5 mM cyclohexanone was added to the vials to induce 1-undecene synthesis, and the vials were further incubated for 23 hours at 30 °C and 300 rpm. The liquid culture samples from *M. tundripaludum* SV96 and *M. rosea* SV97 cultivation without *A. baylyi* ADP1 *‘tesA-undA* were included as the experimental control. The growth of *A. baylyi* ADP1 ‘*tesA-undA* was estimated based on difference of OD_600nm_ between the tests and controls at 24-h incubation (ΔOD_600nm_).

### 2.5. Analytical methods

Liquid samples were filtered through a 0.2 μm membrane (Chromafil® Xtra PET-20/25, Macherey-Nagel, Germany) prior to the analysis of liquid metabolites using a Shimadzu high-performance liquid chromatograph (HPLC) equipped with Rezex RHM-Monosaccharide H^+^ column (Phenomenex, USA) as described in Okonkwo et al. (2018). The gas samples in the headspace (CH_4_, CO_2_, O_2_ and N_2_) were measured using a Shimadzu gas chromatograph GC-2014 equipped with a thermal conductivity detector (TCD) and a Carboxen-1000 60/80 column (Agilent Technologies, USA). The oven temperature was held at 35 °C for 3.75 min, increased 30 °C min^−1^ to 150 °C for 3 min. The injector and detector were 155 °C and 160 °C, respectively. Helium was used as a carrier gas at 30 mL min^−1^.

The production of 1-undecene in headspace was detected using gas chromatography-mass spectrometry (GC-MS) as described by Luo et al. (2019). Compounds were identified using the NIST/EPA/NIH Mass Spectral Library (NIST 05). The bacterial growth was measured as OD_600nm_ using an Ultrospec 500 pro spectrophotometer (Amersham Biosciences, UK) and as cell dry weight (CDW) using gravimetric method. The conversion factor between CDW and OD_600nm_ obtained from the experimental measurement was 0.2914 g L^−1^ OD^−1^ (R^2^ = 0.9948) and 0.3015 g L^−1^ OD^−1^ (R^2^ = 0.9892) for *M. tundripaludum* SV96 and *M. rosea* SV97, respectively.

Wax ester production was detected by using lipid extraction and thin layer chromatography (TLC) analysis. Lipid extraction was done by using methanol-chloroform extraction described by Santala et al. (2011) and the obtained lipid phase layer was used for TLC analysis to visualize the total lipid composition. TLC analysis was done as described by Santala et al. (2011). Briefly, 60 μl of lipid phase of a sample was applied to TLC plate (10 × 20 cm HPTLC Silica Gel 60 F254 glass plates with 2.5 × 10 cm concentrating zone, Merck, USA) using n-hexane:diethyl ether:acetic acid of 90:15:1 as a mobile phase. The visualization was done by dyeing with iodine and Jojoba oil was used as a standard.

### 2.6. Statistical analysis

The statistical analysis was performed using Minitab16.0 (USA). The significant differences of the obtained data sets (e.g., growth of the tested strains, gas concentrations and utilization, concentrations and yields of the products produced by each strain) within the varied treatment were analyzed using one-way of variance (ANOVA) with Tukey’s multiple comparison tests at the 95% confidence interval, where *P-value* ≤ 0.05 was considered statistically significant.

## 3. Results and discussion

### 3.1. CH_4_-derived organic acid production of *M. tundripaludum* SV96 and *M. rosea* SV97 in different gas supplementation tests

Both *M. tundripaludum* SV96 and *M. rosea* SV97 used CH_4_ as the sole carbon and energy sources and O_2_ as an electron acceptor. In all gas supplementation tests, *M. tundripaludum* SV96 showed higher in biomass production than *M. rosea* SV97 (Fig. 3). Highest biomass production of both methanotrophs was observed in the supplementation of both CH_4_+air (test I) on day 14 at concentrations of 0.60 ± 0.03 and 0.48 ± 0.05 g CDW L^−1^ for *M. tundripaludum* SV96 and *M. rosea* SV97, respectively. In all tests, the growth yield of *M. rosea* SV97 was lower than *M. tundripaludum* SV96 (*P < 0.05*) due to typical carbon assimilation pathway of Type I methanotrophs (RuMP) showing more efficient channeling of CH_4_-carbon to biomass than in Type II methanotrophs (serine cycle) (Kalyuzhnaya et al., 2015).

**Fig. 3.**
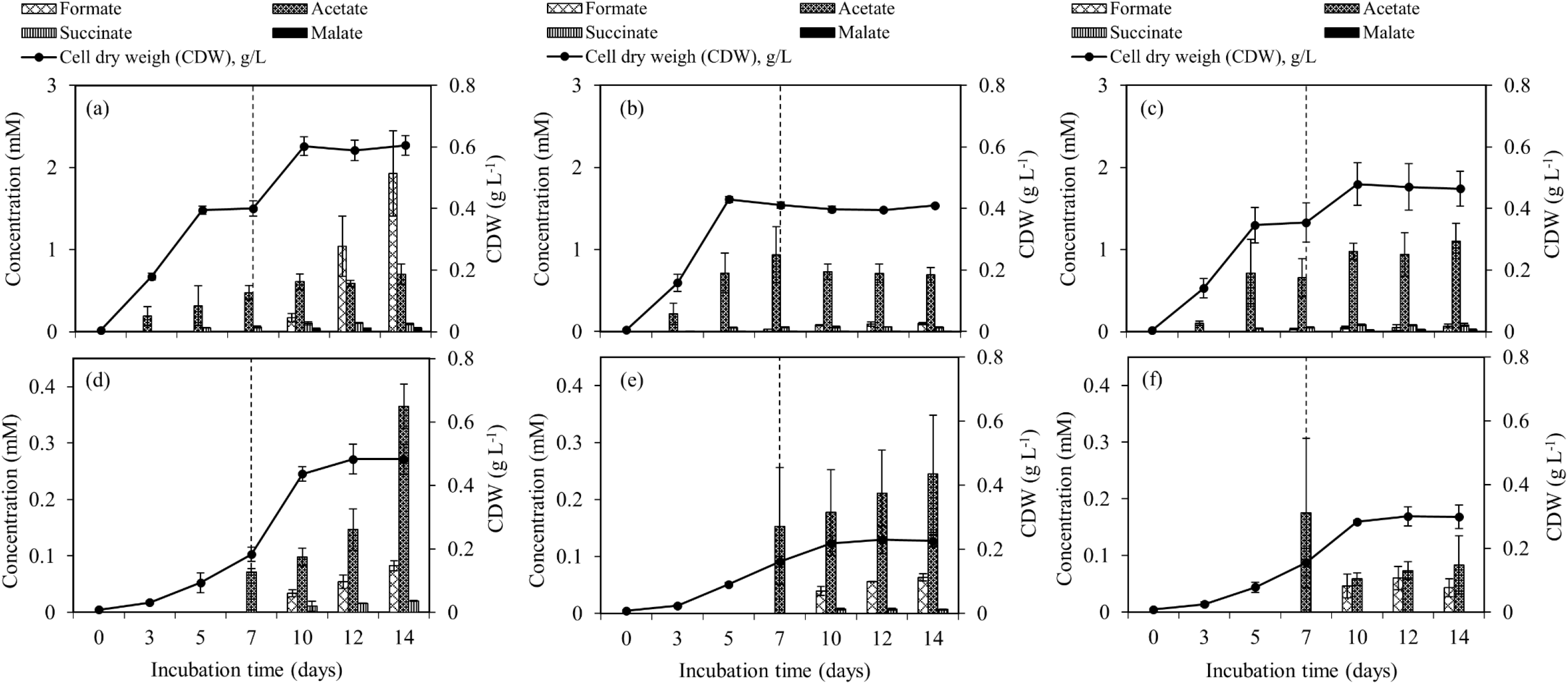
The accumulation of organic acids production and biomass production during the 14-day incubation of *M. tundripaludum* SV96 (a-c) and *M. rosea* SV97 (d-f) under three different gas supplementation schemes applied on day 7: (i) both CH_4_ and air added (a and d), (ii) only CH_4_ added (b and e), and (iii) only air added (c and f). Error bars indicate the standard deviation of triplicate samples.

Spent media of both methanotrophs contained similar organic acid compounds including formate, acetate, and succinate, while malate was only present in the spent medium of *M. tundripaludum* SV96. However, the results suggested that *M. tundripaludum* SV96 was more efficient and promising in CH_4_ conversion into organic acids than *M. rosea* SV97. The spent medium of *M. tundripaludum* SV96 contained higher concentrations and yields of organic acids than those in *M. rosea* SV97 in all gas supplementation tests (*P < 0.05*) (Fig. 3). The efficiency of conversion of CH_4_-C into organic acids was 4.8-7.0% and 0.7-1.8% (of consumed CH_4_-C) for *M. tundripaludum* SV96 and *M. rosea* SV97, respectively (Table S1). This difference could be due to their different carbon assimilation pathways, i.e. that RuMP pathway of Type I methanotrophs efficiently links to glycolytic pathway where pyruvate is converted to organic acids, while the serine cycle of Type II methanotrophs has high flux through acetyl-CoA (Kalyuzhnaya et al., 2015) which is derived into an intracellular storage compound, like polyhydroxybutyrate (PHB), as previously observed in *M. rosea* SV97 during isolation characterization (Wartiainen et al., 2006b). Regarding the production of organic acids in Type II methanotrophs, which has not been widely reported, Costa et al. (2000) observed methanotroph-driven CH_4_ conversion into acetate which was subsequently consumed by heterotrophs in the denitrification bioreactor using CH_4_ as an electron donor. Vecherskaya et al. (2009) also observed succinate, acetate and 2,3-butanediol excreted in culture medium of Type II methanotroph, *Methylocystis parvus*, during CH_4_ oxidation under microaerobic and anaerobic conditions. Based on ^13^C analysis in their study, those organic acids were likely from PHB degradation during microaerobic conditions (5-10% O_2_) (Vecherskaya et al. 2009).

During 14-day incubation in all gas supplementation tests, organic acid concentrations gradually accumulated corresponding with the increase of biomass concentration in both methanotroph liquid cultures (Fig. 3). On days 10-14 onwards, different gas supplementation tests likely induced different conditions to headspace including microaerobic (O_2_-limited), anaerobic and aerobic conditions. These conditions resulted in O_2_/CH_4_ molar ratios in headspace of 0.2-0.3, < 0.01 and 2-10 for the addition of CH_4_ + air, only CH_4_ and only air, respectively (Fig. S1). Interestingly, the three different gas supplementation tests in our study showed similar total organic acid production yields (Fig. 4a and b). Lee et al. (2021) compared the production of organic acids in *M. capsulatus* Bath culture under various conditions including aerobic, oxygen-limited, sulfur-limited and nitrogen-limited conditions. The authors observed acetate production in all studied conditions, and nitrate-nitrogen limitation induced the highest acetate production (approximately 1.9 mmol-acetate g^−1^ CDW). Furthermore, the previous studies on organic acid production by type I methanotrophs (Lee et al., 2021; Takeuchi and Yoshioka, 2021) reported that acetate and formate were also produced under aerobic conditions but in lower concentrations than in O_2_-limited conditions. In our study, however, the three different gas supplementation tests likely varied in the distribution of organic acids contained in the spent medium, particularly, of *M. tundripaludum* SV96, type I methanotrophs. Regarding organic acid production yields per consumed CH_4_-carbon, both formate (3.5%) and acetate (2.5%) were dominant in the test with CH_4_+air supplementation (test I), while only acetate (5.1%) was dominant in the test with only air supplementation (test III) (Fig. 4c and d). These results suggest that it would be possible to target dominant organic acid compounds by controlled feeding CH_4_ and O_2_. The excretion of high formate concentration of *M. tundripaludum* SV96 at CH_4_+air supplementation (test I) might be due to imbalanced growth during O_2_-limited conditions, which was previously reported in type I methanotrophs by Gilman et al. (2017). This circumstance also corresponded with pH reduction observed only in the test with CH_4_+air supplementation (test I) (Fig. S2).

**Fig. 4.**
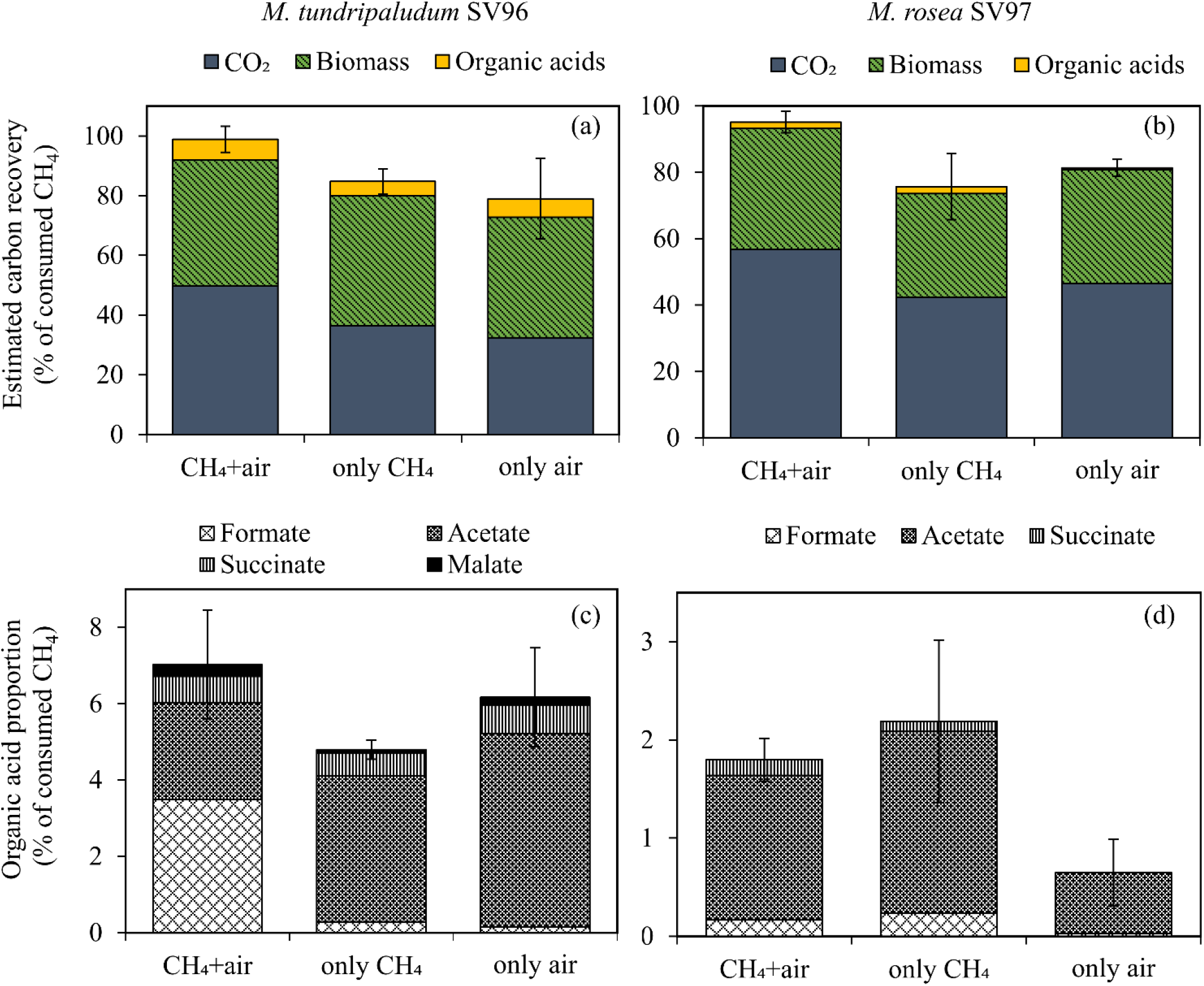
Carbon mass balance applied to CH_4_ oxidation to CO_2_, biomass, and organic acids (a and b) and distribution of each organic acid compound (c, d) of *M. tundripaludum* SV96 (left) and *M. rosea* SV97 (right) in three different gas supplementation tests for the 14-day incubation. Error bars indicate the standard deviation of sum of total yields in triplicate.

Comparing to organic acid production in different Type I methanotroph species under O_2_-limited conditions, *M. tundripaludum* SV96 used in our study showed comparable and higher organic acid yields per biomass production (CDW), i.e., for formate, acetate, and succinate (Table 1). Those previous studies of organic acid production from microaerobic CH_4_ oxidation cultivated the methanotrophs with the initial headspace CH_4_ and O_2_ concentrations of 20% and 5%, respectively (Gilman et al., 2017, 2015; Kalyuzhnaya et al., 2013). However, in this study, *M. tundripaludum* SV96 was prior cultivated aerobically before allowing O_2_ limiting condition to occur. Whether the prior cultivation in aerobic conditions enhanced organic acid production in the subsequent microaerobic conditions requires further studies. Additionally, the organic acid yields of *M. tundripaludum* SV96 obtained from our study (4.8-7.0% of consumed CH_4_-carbon) were higher than those of *M. capsulatus* Bath (<5% of consumed CH_4_) (Lee et al., 2021), while *M. alcaliphilum* 20Z enabled to convert 40-50% of the consumed CH_4_ into mostly acetate and formate under O_2_-limited conditions (20% CH_4_:5% O_2_) (Kalyuzhnaya et al., 2013).

**Table 1.**
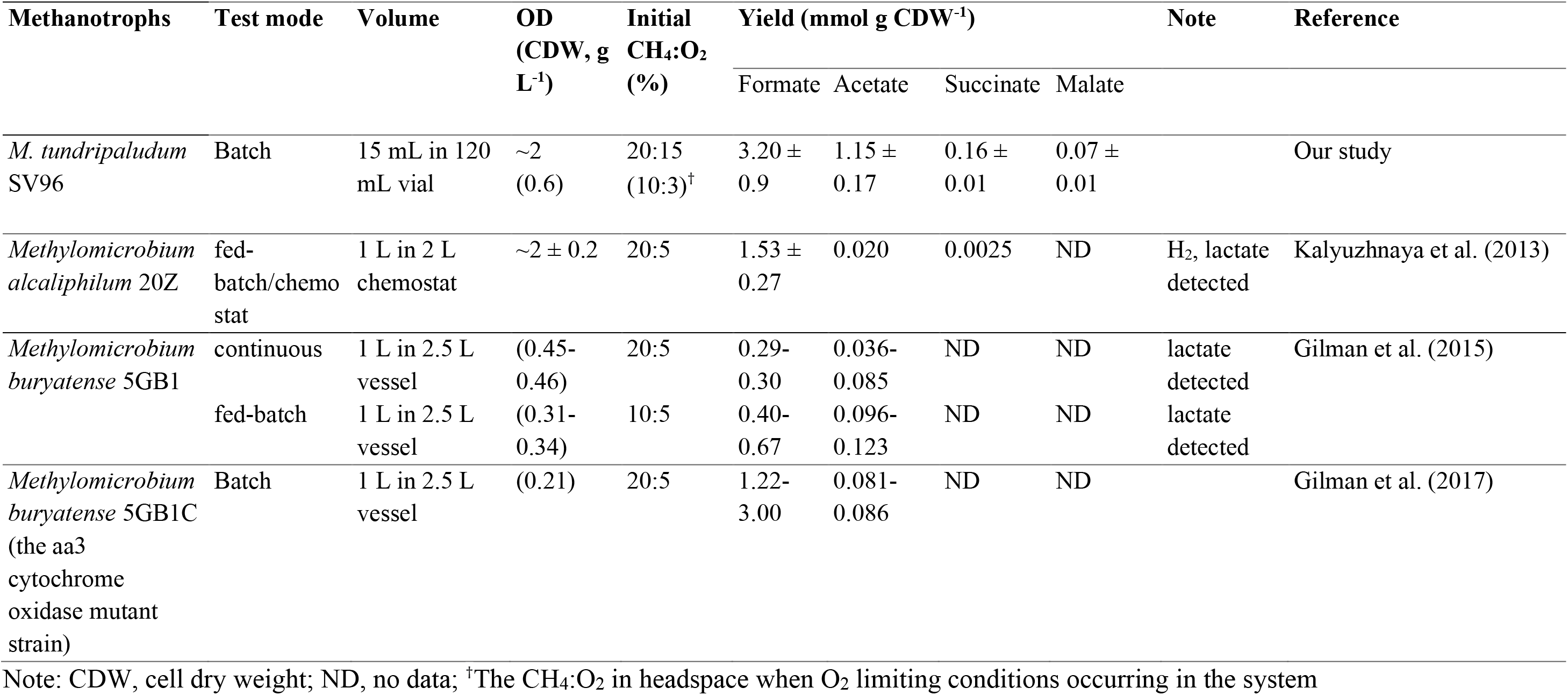
The production of organic acids and other metabolites obtained from different Type I methanotrophs cultivated under O_2_ limiting conditions.

### 3.2. Utilization of organic acids rich spent medium of methanotrophs by *A. baylyi* ADP1

Wild type *A. baylyi* ADP1 cells enabled to grow in the organic acid-rich spent media of both methanotrophs, containing formate, acetate, and succinate (and malate for *M. tundripaludum* SV96). During 4-h incubation, *A. baylyi* ADP1 cells incubated in the spent media from *M. tundripaludum* SV96 and *M. rosea* SV97 grew to an OD_600nm_ of 0.14 ± 0.01 and 0.12 ± 0.03, respectively (Fig. S3). Except for formate, acetate, succinate, and malate were utilized as carbon sources for biomass assimilation (Fig. S3). *A. baylyi* ADP1 cells do not utilize formate as a sole carbon source, but it is rather used for maintaining the cellular redox balance (Kannisto et al., 2015). The growth of *A. baylyi* ADP1 was also confirmed by the present of fatty acid fractions derived from *A. baylyi* ADP1 biomass on TLC plates. Typically, *A. baylyi* ADP1 produce wax esters as carbon storage compounds which often associated with growth (Mangayil et al., 2019). However, they were not detected in this study (Fig. S4). This could be due to rapid degradation of the accumulated wax ester during carbon limiting conditions. Under carbon limiting conditions *A. baylyi* ADP1 cells utilize the stored carbon for maintaining cellular activities (Fixter et al., 1986; Salmela et al., 2018). The application of using the spent media of methanotrophs for wax production would be possible for further study but likely require higher quantity of organic acids.

### 3.3. 1-alkene synthesis from organic acids rich spent medium of methanotrophs

After confirming the growth of wild type *A. baylyi* ADP1 on the methanotroph spent media, the organic acid-rich spent media were hereby used as carbon sources for 1-alkene production by an engineered *A. baylyi* ADP1 ‘*tesA-undA*. The results showed that *A. baylyi* ADP1 ‘*tesA-undA* completely utilized acetate (0.3 mM), succinate (0.05 mM), and malate (0.2 mM) present in the spent media of methanotrophs, except formate, similar to wild type ADP1 cells (Fig. 5a). The production of 1-undecene from *M. tundripaludum* SV96 spent medium (14.1 ± 2.7 μg L^−1^) was higher than that from *M. rosea* SV97 cultivations (1.0 ± 0.5 μg L^−1^) (Fig. 5c), corroborating with the organic acid concentrations (Fig. 5a and 5b). Likewise, the growth of *A. baylyi* ADP1 ‘*tesA-undA* was higher in spent medium from *M. tundripaludum* SV96 (0.115 ± 0.06 OD_600nm_) than from *M. rosea* SV97 (0.070 ± 0.054 OD_600nm_) (Fig. 5c). The production of 1-undecene was not detected in the control cultivations (Fig. S5). In addition, the growth occurred during the co-cultivation was from solely *A. baylyi* ADP1 ‘*tesA-undA* as both methanotrophs could not grow without CH_4_ as a carbon source.

**Fig. 5.**
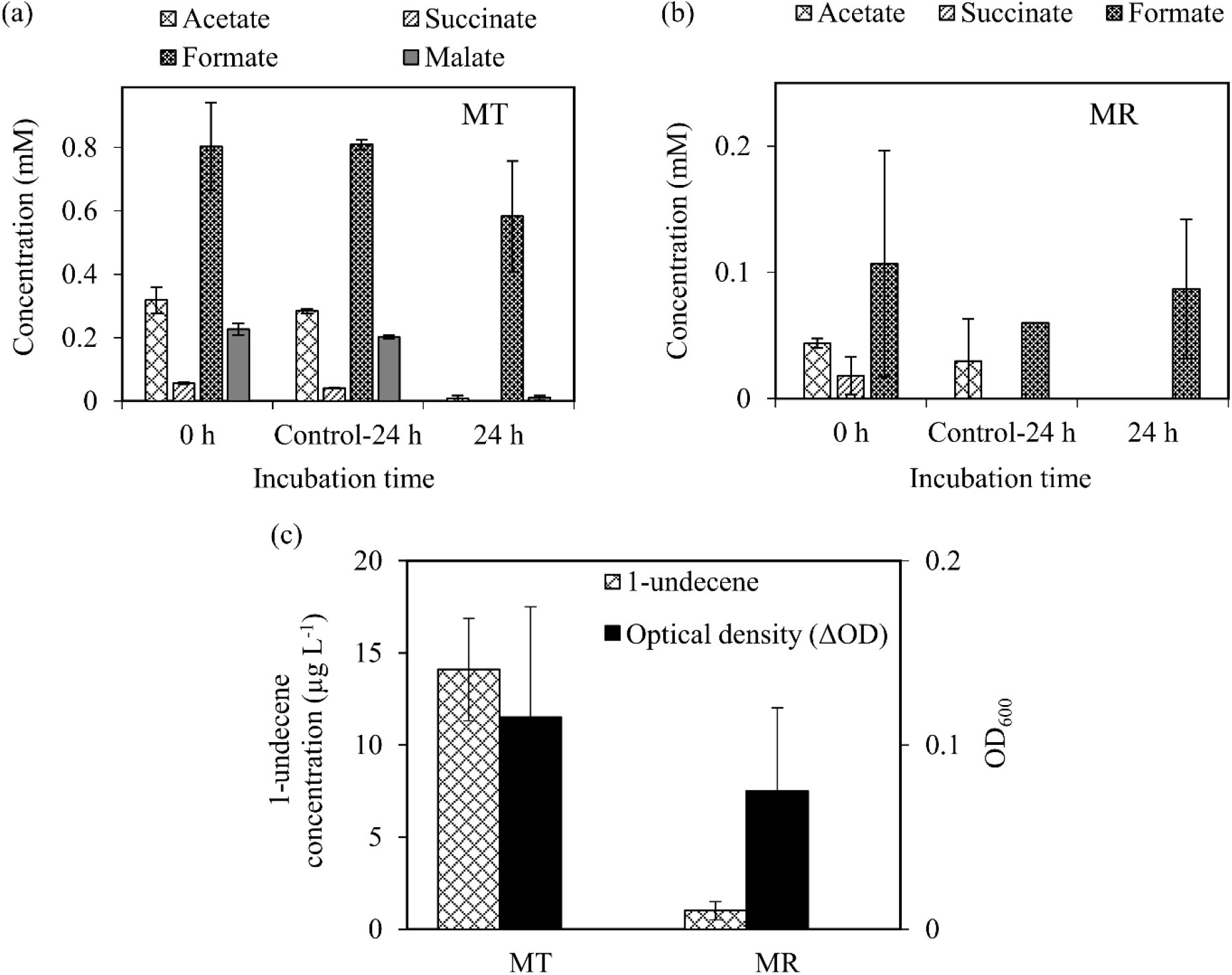
Organic acids utilization by *A. baylyi* ADP1 *‘tesA*-*undA* in spent media obtained from the cultivation of *M. tundripaludum* SV96 (MT) (a) and *M. rosea* SV97 (MR) (b) under O_2_ limiting conditions. Growth and 1-undecene production from *A. baylyi* ADP1 ‘*tesA*-*undA* after 24 h incubation (c). ΔOD indicates the difference in OD_600nm_ of *A. baylyi* ADP1 ‘*tesA*-*undA* between 0 and 24 h. Error bars indicate the standard deviation of four replicate samples. The spent media from methanotrophs fermentation without *A. baylyi* ADP1 were used as control (Control-24 h).

The obtained 1-undecene concentrations in this study (1.0 and 14.1 μg L^−1^) were lower than that previously reported for *A. baylyi* ADP1 ‘*tesA-undA* (Luo et al., 2019; Salmela et al., 2020). This phenomenon can be attributed towards the higher substrate concentrations used in those studies. For example, Luo et al. (2019) developed a high ferulate-tolerant *A. baylyi* ADP1 strain for 1-undecene production using adaptive laboratory evolution. The authors tested the use of high concentration of ferulate (100 mM) as a sole carbon source and obtained the 1-undecene production titer of 72 ± 7.5 μg L^−1^ (yield of 1.0 μmol mol C_substrate_^−1^). In another study, Salmela et al. (2020) obtained 1-undecene concentration of up to ~107 ± 8 μg L^−1^ from a two-stage system for 1-undecene production from cellulose which was converted into organic metabolites by *Clostridium cellulolyticum* (containing 5.2 mM glucose, 4.9 mM acetate and 6.8 mM lactate). The authors reported the highest 1-undecene production yield of ~35 μmol mol C_substrate_^−1^ using lactate as a substrate. In our study, 1-undecene production from the spent media of methanotrophs was promising and comparable to those in previous studies, resulting in 1-undecene yields of 68.9 ± 11.6 and 40.6 ± 19.8 μmol mol C_substrate_^−1^ for *M. tundripaludum* SV96 and *M. rosea* SV97, respectively. The obtained carbon recovery for 1-undecene production accounted for 0.065% and 0.045% of consumed total organic acids-carbon for *M. tundripaludum* SV96 and *M. rosea* SV97, respectively (Table S2). The spent media of methanotrophs likely did not contain intermediate compounds being toxic to growth and 1-undecene production of *A. baylyi* ADP1 ‘*tesA-undA*. Furthermore, the media could be directly used for the cultivation without purification or additional downstream processing.

The results indicate that the spent media from microaerobic fermentation by methanotrophs is an excellent carbon source for heterotrophs. This two-stage bacterial process extended the range of CH_4_-derived products to C11 compound (1-undecene). In further studies, the process optimization, scale-up as well as statistical optimization of the system for methanotroph cultivation will be developed to enhance organic acid concentration, which will consequently improve the production efficiency of *A. baylyi* ADP1. As produced organic acids, particularly acetate and succinate, by methanotrophs were also observed under aerobic conditions, this study can also lead to develop a single system for CH_4_ conversion into high value-added products using coculture of methanotrophs and *A. baylyi* ADP1.

## 4. Conclusion

A two-step bioprocess setup was designed for successful integration of microaerobic methane fermentation with aerobic synthesis. This study shows a proof of concept for integrating greenhouse gas (GHG) utilization and platform chemical production: the application of organic acids produced by methanotrophs for 1-undecene (C11) production. A Type I methanotroph, *M. tundripaludum* SV96, showed higher potential for organic acid production than a Type II methanotroph, *M. rosea* SV97 under aerobic and microaerobic conditions. The organic acids-rich spent media of methanotrophs could be directly used as a medium for the cultivation of the wild type *A. baylyi* ADP1 and the engineered *A. baylyi* ADP1 ‘*tesA-undA* without additional downstream processes or purification. Acetate, succinate, and malate contained in the spent media were completely utilized by *A. baylyi* ADP1 ‘*tesA-undA* for their 1-undecene production. The highest yield of 1-undecene was obtained from spent medium of *M. tundripaludum* SV96 at 68.9 ± 11.6 μmol mol C_substrate_^−1^.

## Acknowledgement

The research work was supported by Kone foundation [grant number 201803224] and Academy of Finland [grant number 323214]. The authors would like to acknowledge Jin Luo for helping with using GC-MS for 1-undecene measurement.

## Appendices

Supplementary data is provided as a separate file attached with the manuscript.

