## Supplementary data for "Batch experiments demonstrating a two-stage bacterial process coupling methanotrophic and heterotrophic bacteria for 1-alkene production from methane"

**Table S1.** The carbon mass balance and distribution of consumed $CH_4$-carbon after 14-day incubation of the experiments of *M. tundripaludum* SV96 (MT) and *M. rosea* SV97 (MR). All tests were incubated with both $CH_4$ and air (20% $CH_4$ and 80% air) in headspace on day 0 and three different gas supplementation schemes applied on day 7.

| | | | *M. tundripaludum* SV96 | | | *M. rosea* SV97 | | |
|---|---|---|---|---|---|---|---|---|
| Tests | | | **I: $CH_4$ + air** | **II: only $CH_4$** | **III: only air** | **I: $CH_4$ + air** | **II: only $CH_4$** | **III: only air** |
| Carbon mass (μmol) | Consumed $CH_4$ | | 552.21 | 358.73 | 438.90 | 498.29 | 266.24 | 324.32 |
| | Produced $CO_2$ | | 275.13 | 130.20 | 141.56 | 281.89 | 112.38 | 150.81 |
| | Biomass[†] | | 231.91 | 156.17 | 177.27 | 182.44 | 83.04 | 111.10 |
| | Produced organic acids | Formate(C1) | 19.28 | 0.99 | 0.67 | 0.83 | 0.63 | 0.11 |
| | | Acetate (C2) | 14.00 | 13.80 | 22.09 | 7.30 | 4.90 | 2.10 |
| | | Succinate (C4) | 3.79 | 2.12 | 3.26 | 0.80 | 0.27 | n.d. |
| | | Malate (C4) | 1.69 | 0.31 | 0.93 | n.d.[‡] | n.d. | n.d. |
| | | Total organic acids | 38.77 | 17.22 | 26.95 | 8.93 | 5.81 | 1.50 |
| | Total carbon products | | 545.80 ± 13.44 | 303.59 ± 8.61 | 345.79 ± 51.45 | 473.26 ± 23.74 | 201.22 ± 22.62 | 263.42 ± 28.30 |
| Carbon distribution of consumed $CH_4$ (%) | Produced $CO_2$ | | 49.8 | 36.4 | 32.4 | 56.7 | 42.3 | 46.5 |
| | Biomass | | 42.0 | 43.6 | 40.4 | 36.6 | 31.2 | 34.2 |
| | Produced organic acids | Formate(C1) | 3.5 | 0.3 | 0.2 | 0.2 | 0.2 | 0.0 |
| | | Acetate, (C2) | 2.5 | 3.8 | 5.1 | 1.5 | 1.8 | 0.6 |
| | | Succinate (C4) | 0.7 | 0.6 | 0.7 | 0.2 | 0.1 | 0.0 |
| | | Malate (C4) | 0.3 | 0.1 | 0.2 | 0.0 | 0.0 | 0.0 |
| | | Total organic acids | 7.0 | 4.8 | 6.2 | 1.8 | 2.2 | 0.7 |
| Carbon recovery (%) | | | 98.9 ± 4.4 | 84.7 ± 4.2 | 79.0 ± 13.5 | 95.1 ± 3.2 | 75.7 ± 10.0 | 81.1 ± 2.6 |
| Organic acid yield (mmol $g^{-1}$ CDW) | | Formate(C1) | 3.19 ± 0.89 | 0.24 ± 0.03 | 0.15 ± 0.07 | 0.17 ± 0.02 | 0.28 ± 0.04 | 0.13 ± 0.05 |
| | | Acetate, (C2) | 1.15 ± 0.17 | 1.68 ± 0.20 | 2.41 ± 0.60 | 0.76 ± 0.14 | 1.07 ± 0.35 | 0.28 ± 0.27 |
| | | Succinate (C4) | 0.16 ± 0.01 | 0.13 ± 0.01 | 0.17 ± 0.02 | 0.04 ± 0.00 | 0.03 ± 0.00 | n.d. |
| | | Malate (C4) | 0.07 ± 0.01 | 0.02 ± 0.00 | 0.05 ± 0.01 | n.d. | n.d. | n.d. |

Note: [†]Chemical formula for biomass is from *Methylococcus capsulatus* = $CH_2O_{0.5}N_{0.27}$ (25.78 g $mol^{-1}$) (Popovic, 2019); [‡] n.d., not detected

Reference: Popovic, M. (2019). Thermodynamic properties of microorganisms: determination and analysis of enthalpy, entropy, and Gibbs free energy of biomass, cells and colonies of 32 microorganism species. *Heliyon*, *5*, e01950. https://doi.org/10.1016/j.heliyon.2019.e01950

**Table S2.** Carbon mass balance applied to organic acid production by methanotrophs cultivated in a 500 mL vial and 1-undecene production from organic acid-rich spent medium of methanotrophs by *A. baylyi* ADP1 *tesA-undA* cultivated in a 5 mL vial.

| | | MT | MR | MT | MR |
|---|---|---|---|---|---|
| | | Mass | | Carbon mass | |
| **Carbon balance of organic acids production by MOBs (in 500 mL vial)** | **Substrate consumed by by MOBs** | | | | |
| | | µmol | | µmol | |
| | $CH_4$ consumed by methanotrophs | 5218.43 | 5536.87 | 5218.43 | 5536.87 |
| | **Products** | | | | |
| | Produced $CO_2$ | 1962 | 1668 | 1962 | 1668 |
| | Produced biomass | 2261 | 3275 | 2261 | 3275 |
| | Formate ($CHO_2^-$) | 80.32 | 10.68 | 80.32 | 10.68 |
| | Acetate ($C_2H_3O_2^-$) | 31.87 | 4.38 | 63.75 | 8.77 |
| | Succinate ($C_4H_4O_4^{2-}$) | 5.58 | 1.81 | 22.32 | 7.26 |
| | Malate ($C_4H_4O_5^{2-}$) | 22.66 | 0.00 | 90.65 | 0 |
| | Total | | | 4480.0 | 4969.3 |
| | | | | % Carbon recovery | |
| | Recovery of C and e- for organic acids production from CH4 | | | 4.93 | 0.48 |
| | Recovery of C and e- for CO2 production from CH4 | | | 37.60 | 30.12 |
| | Recovery of C and e- for biomass production from CH4 | | | 43.32 | 59.14 |
| | %Total carbon & electron recovery | | | 85.85 | 89.75 |
| **Carbon balance of 1-undecene production from organic acids by ADP1 (in 5 mL vial)** | **Substrate consumed by *A. baylyi tesA-undA*** | | | | |
| | | µmol | | µmol | |
| | Acetate ($C_2H_3O_2^-$) | 1.55 | 0.22 | 2.32 | 0.44 |
| | Succinate ($C_4H_4O_4^{2-}$) | 0.28 | 0.09 | 1.12 | 0.36 |
| | Malate ($C_4H_4O_5^{2-}$) | 1.08 | - | 4.32 | - |
| | Total | | | 7.76 | 0.80 |
| | **Product** | | | | |
| | 1-undecene ($C_{11}H_{22}$) | 0.0005 | 0.000033 | 0.00503 | 0.00036 |
| | 1-undecene yield (µmol $mol^{-1}$ Carbon substrate) | 58.86 | 40.63 | | |
| | | | | % Carbon recovery | |
| | Carbon recovery for 1-undecene production from organic acids | | | 0.065 | 0.045 |
| | Carbon recovery for 1-undecene production from consumed $CH_4$ | | | 0.0001 | 0.00001 |

**Gas composition during 14-day incubation of methanotrophs in this study**

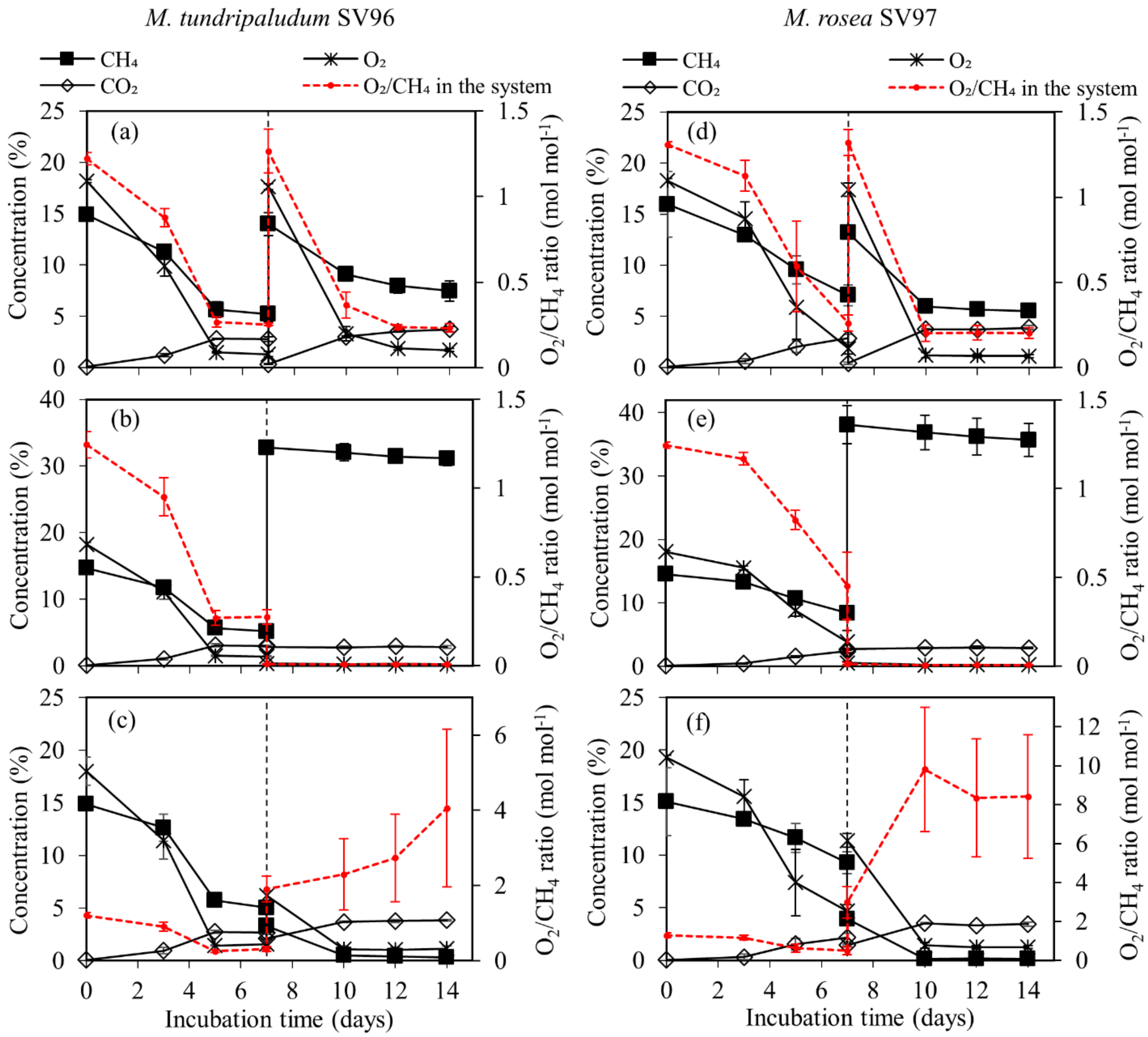


**Fig. S1.** Gas compositions ($CH_4$, $O_2$ and $CO_2$) in headspace during the 14-day incubation of *M. tundripaludum* SV96 (left column) and *M. rosea* SV97 (right column) under three different gas supplementation schemes applied on day 7: (i) both $CH_4$ and air added (a,d), (ii) only $CH_4$ added (b,e), and (iii) only air added (c,f). Error bars indicate the standard deviation of triplicate samples.

### Profiles of pH and optical density (OD) during cultivation of methanotrophs

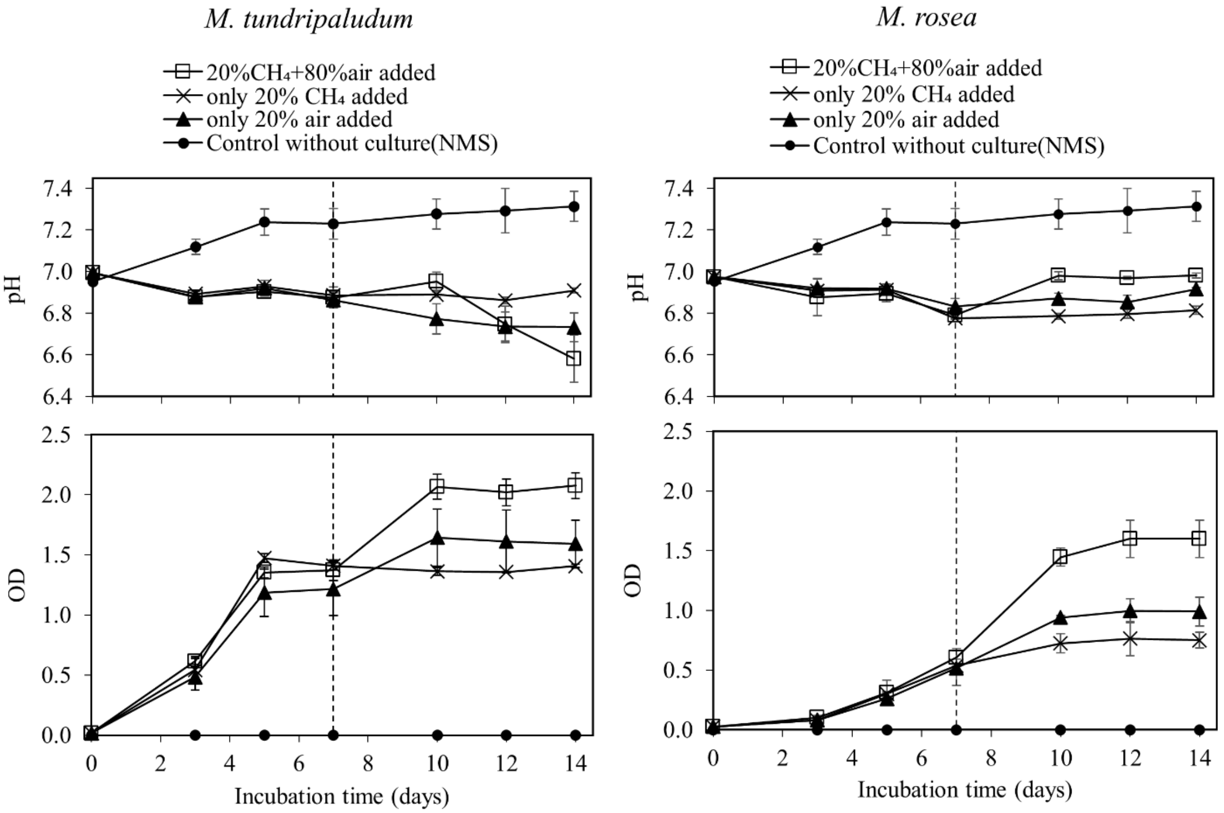


**Fig. S2.** Profiles of pH and OD during 14-day incubation of *M. tundripaludum* SV96 (a) and *M. rosea* SV97 (b) under three different gas supplementation schemes: (test I) both $CH_4$ and air added, (test II) only air added, and (test III) only $CH_4$ added. The error bars represent the standard deviations of triplicate samples.

### The growth of wild type *A. baylyi* ADP1 on spent medium of methanotrophs

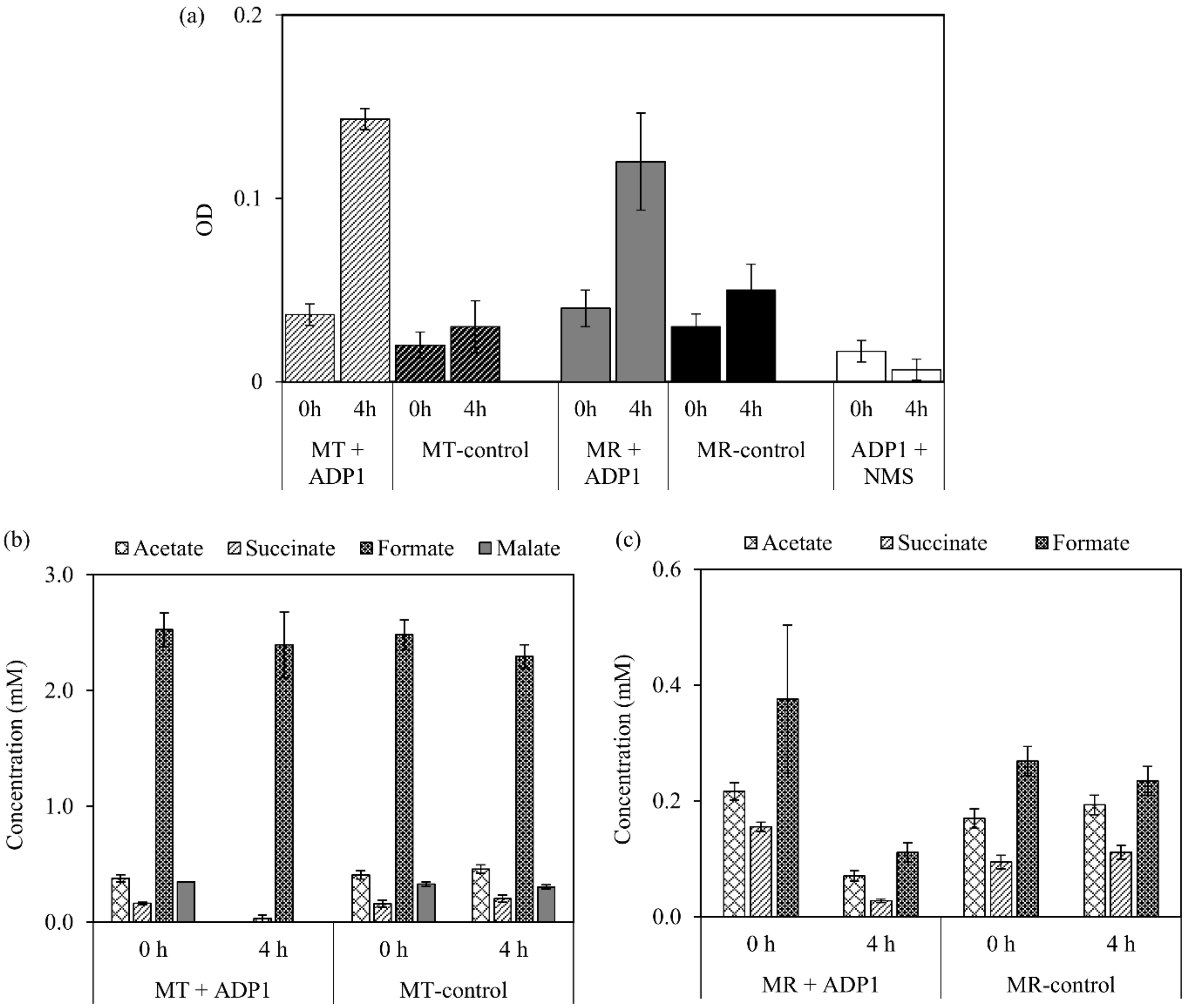


**Fig. S3.** Growth of wild type *A. baylyi* ADP1 after 4-h cultivation in the methanotroph spent media (a) and concentrations of organic acids contained in the spent media of *M. tundripaludum* SV96 (MT) (b) and *M. rosea* SV97 (MR) (c). Error bars indicate the standard deviation of duplicate samples. The incubations of spent media of MT and MR without *A. baylyi* ADP1 (MT- and MR-controls) and *A. baylyi* ADP1 with NMS fresh medium (ADP1 + NMS) were used as controls.

**Visualization of total lipid composition**

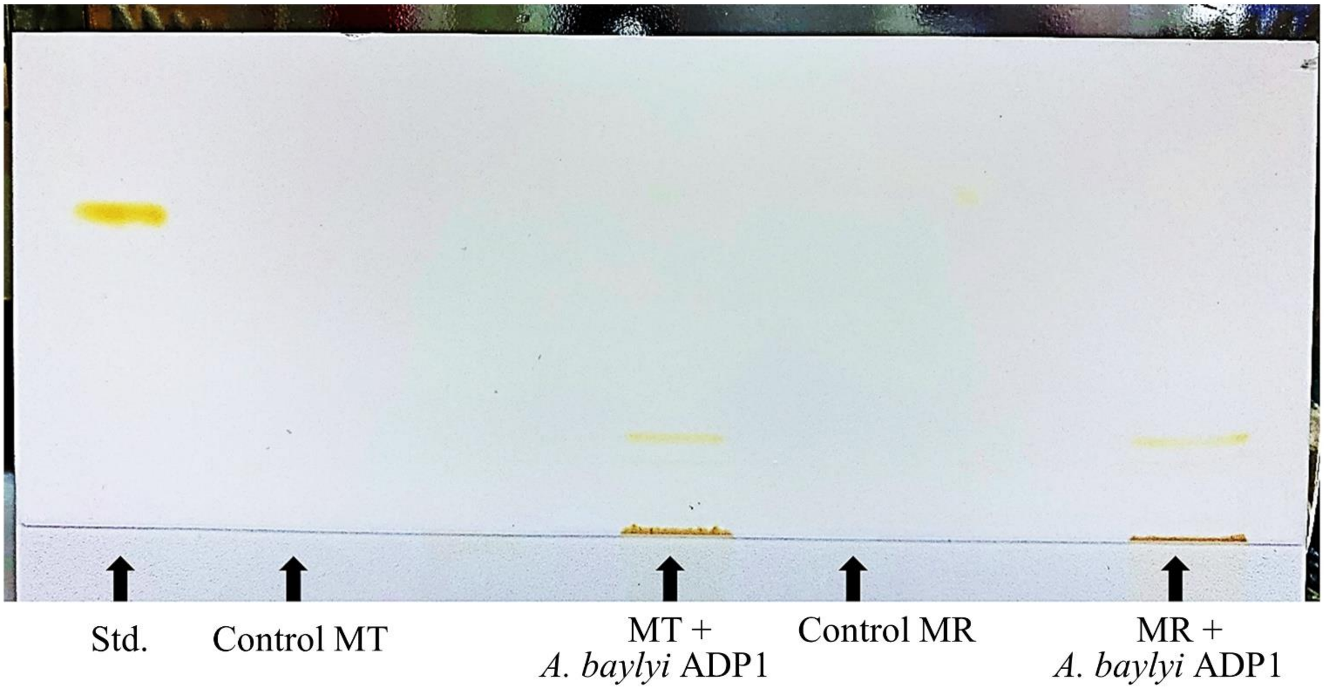


**Fig. S4.** The volumetric WE production in *A. baylyi* ADP1 cultivated in the spent media of *M. tundripaludum* SV96 (MT) and *M. rosea* SV97 (MR) were determined by thin layer chromatography analysis. Jojoba oil was used as the standard (Std.). The spent media without *A. baylyi* ADP1 were used as control1 (MT-control and MR-control).

### The present of 1-undecene in the cultivation detected by GC-MS

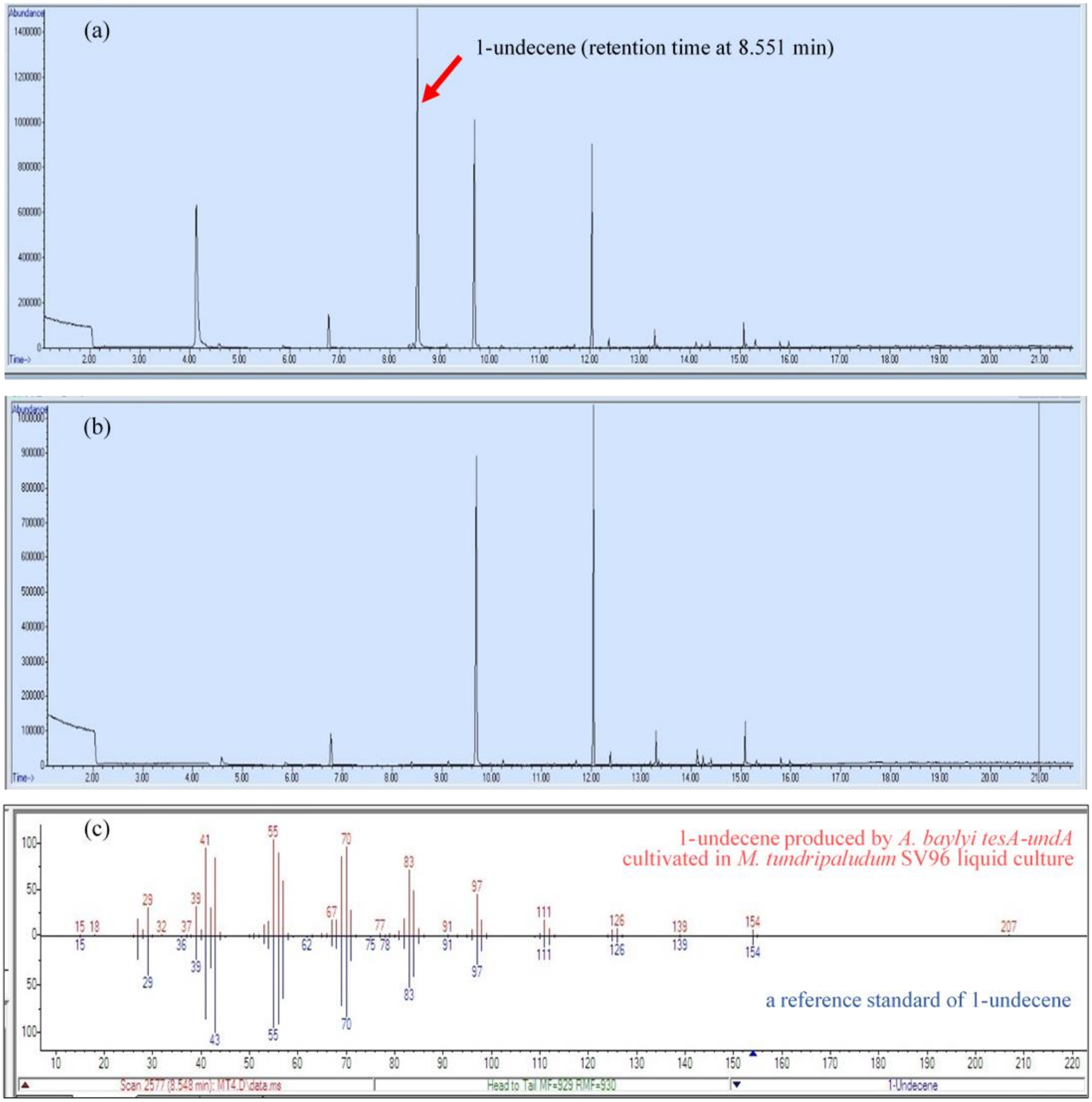


**Fig. S5.** GC-MS analysis for identifying 1-undecene in headspace of vials cultivating *A. baylyi tesA-undA* in *M. tundripaludum* SV96 spent medium (a) and the control cultivation of *M. tundripaludum* SV96 without *A. baylyi tesA-undA* (b). The reverse panel shows the reference standard of 1-undecene from NIST mass spectrometry data center (in blue) compared to the produced 1-undecene in the culture supernatant (in red) (c).
